# Applicability of the mutation-selection balance model to population genetics of heterozygous protein-truncating variants in humans

**DOI:** 10.1101/433961

**Authors:** Donate Weghorn, Daniel J. Balick, Christopher Cassa, Jack Kosmicki, Mark J. Daly, David R. Beier, Shamil R. Sunyaev

## Abstract

The fate of alleles in the human population is believed to be highly affected by the stochastic force of genetic drift. Estimation of the strength of natural selection in humans generally necessitates a careful modeling of drift including complex effects of the population history and structure. Protein truncating variants (PTVs) are expected to evolve under strong purifying selection and to have a relatively high per-gene mutation rate. Thus, it is appealing to model the population genetics of PTVs under a simple deterministic mutation-selection balance, as has been proposed earlier [1]. Here, we investigated the limits of this approximation using both computer simulations and data-driven approaches. Our simulations rely on a model of demographic history estimated from 33,370 individual exomes of the Non-Finnish European subset of the ExAC dataset [2]. Additionally, we compared the African and European subset of the ExAC study and analyzed *de novo* PTVs. We show that the mutation-selection balance model is applicable to the majority of human genes, but not to genes under the weakest selection.

## Introduction

In well-adapted populations, the evolutionary dynamics of genes under purifying selection plays a prominent role. Early models describing this phenomenon were purely deterministic, that is they assumed infinite effective population size [3]. In particular, the discussion centered around the concept of mutation-selection balance, when deleterious variants in the population are replenished by mutation against the constant purge of negative selection. Supported by the advent of large sequencing datasets and computer simulations, however, it became clear that the high amounts of non-silent genetic variation observed in real populations cannot be fully explained by mutation-selection balance [4, 5, 6]. Instead, many mutations are only weakly selected against and many populations cannot be approximated to be infinitely large. Both of these factors emphasize the relative importance of stochastic effects, or genetic drift, compared to mutation and selection. Therefore, deterministic mutation-selection balance is not an adequate description of the evolutionary dynamics of deleterious alleles unless both mutation influx and selection strength are sufficiently high to dominate genetic drift. The full mutation-selection-drift balance has been extensively studied using the diffusion approximation [7]. It is now widely appreciated that in humans the complexities of demographic history and changing population size must be explicitly modeled. This has been incorporated in many recent studies that estimated the intensity of selection [8, 9, 10].

One practically important case when both effects of mutation and selection are strong in comparison to genetic drift is in the dynamics of protein truncating variants (PTVs). Most PTVs within a gene are likely to have identical fitness effects and, therefore, can be analyzed in aggregate. The cumulative number of new truncating mutations per gene is two orders of magnitude higher than the per-site expectation. PTVs are also expected to evolve under a pressure of strong negative selection. It is thus appealing to apply a deterministic approximation to model the population genetics of PTVs, as has been proposed by us earlier [1].

Here, we investigated the utility and limits of the deterministic approximation when estimating selection against heterozygous PTVs, *s*_het_, using simulations that explicitly incorporate drift. Further, we studied the effects of population stratification by comparing PTV allele frequencies between the African and non-Finnish European subpopulations of the ExAC dataset. Lastly, we analyzed the estimates of selection strength vis-à-vis the fraction of *de novo* mutations in a recent pedigree sequencing dataset, a direct measure of *s*_het_. Beyond the analysis of the influence of drift on the *s*_het_ estimates, we also present a comparison of *ŝ*_het_ with another measure of protein constraint, pLI [2].

## Impact of genetic drift on PTV allele count

We first analyzed how genetic drift affects the observed number of PTV mutations on a gene, *k*, and its variance in the population. Given an exome sequencing cohort, *k* is the sum of protein-truncating mutations over all sampled chromosomes in the cohort, *n*. The expected value of *k*, E[*k*], is determined by selection as well as by the local cumulative mutation rate, *U*. Provided PTVs are not nearly recessive, E[*k*] is well approximated by the deterministic expression *nU*/*s*_het_ [1, 11]. If mutation and selection are the dominant evolutionary forces, the approximate Poisson nature of the PTV count implies for the variance Var[*k*] = E[*k*] = *nU*/*s*_het_. Meanwhile, if PTV alleles remain in the population for a substantial period of time and thus become subject to the effects of genetic drift, this will tend to increase Var[*k*]. We can express the total variance of *k* as a sum of Poisson sampling variance and variance due to drift (**Methods**).

The impact of genetic drift shows in the population allele frequency spectrum. For given *U* and *s*_het_, this describes the distribution of the gene-specific cumulative frequency of PTVs in the population, *X*, and depends on the demographic history of the population. To gain intuition, we first considered a classic approximation for the allele frequency spectrum under the assumption of strong purifying selection [12]. At equilibrium, the variance of the cumulative PTV allele count *k* in a sample of size *n* chromosomes is then given by Var[*k*] = *nU*/*s*_het_ (1 + *n*/(4*N*_e_*s*_het_)). Notably, this expression approaches the deterministic approximation of *nU*/*s*_het_ if the size of the sample is much smaller than the product of the effective population size, *N*_e_, and the selection coefficient.

## Modeling recent population expansion

Human populations appear to be far from evolutionary equilibrium, and most populations have undergone very rapid, recent growth [13]. With our focus on rare deleterious PTVs 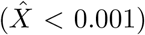, these alleles are expected to be on average young, so the most relevant population size for our purposes is the recent effective population size, corresponding to the lifetime of an average very rare deleterious allele. Current literature estimates of the present day effective size in the European population span the wide range of 0.5 to 8 million individuals [9, 14, 15, 16]. Although the long-term effective size of the human population is smaller than the sample size of the ExAC dataset, this number, driven by ancestral or bottleneck sizes, is less relevant than the epoch of recent growth for very young alleles under relatively strong natural selection.

To determine the deviation from the deterministic regime, we evaluated relevant features of the demographic history under which rare deleterious variants evolve. We focused on the Non-Finnish European (NFE) subset, representing a majority of the samples (N=33,370) in the ExAC data. The NFE subset has a well studied demographic history from a single ancestry. We used features of the Tennessen *et al.* (2012) [9] model of European demography, including the initial size, bottleneck and initial exponential growth phases. For the final exponential growth phase, we matched properties of rare alleles in the ExAC NFE sample to gauge the final effective population size and corresponding growth rate. We simulated a dense range of exponential growth rates in the final demographic period, and matched neutral simulation results of the downsampled site frequency spectrum to the fraction of synonymous singleton alleles observed in the NFE subpopulation. We focused on non-CpG transversions for these purposes, as CpG and even non-CpG transitions are known to exhibit the effects of recurrent mutations in the ExAC sample due to elevated mutation rates relative to non-CpG transversions. For example, the fraction of synonymous singletons for non-CpG transversions in ExAC NFE is 63.3% (95% CI 62.1-64.6%; ExAC All 60.5%), while the fraction of synonymous singletons for CpG transitions in ExAC NFE is 39.8% (39.1-40.7%; ExAC All 23.9%). The analysis of singletons resulted in a demographic history with a recent population size of 4.3 million individuals, which is consistent with estimates based on the same dataset provided in Harpak *et al.* (2016) [15].

In **Figure 1(a)**, we find the coefficient of variation, 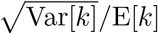, to be in good agreement with the deterministic approximation for genes under strong selection. For *s*_het_ = 0.06, corresponding to the mean of the inferred genome-wide distribution of heterozygous selection coefficients in ref. [1], the inflation of the coefficient of variation in our simulations did not exceed 4%. However, variance indeed diverged from the deterministic approximation for genes under weaker selection (*s*_het_ ≲ 0.02). **Figure 1(a)** also shows that the mean, E[*k*], is unaffected by genetic drift within the analyzed range of parameters.

**Figure 1:**
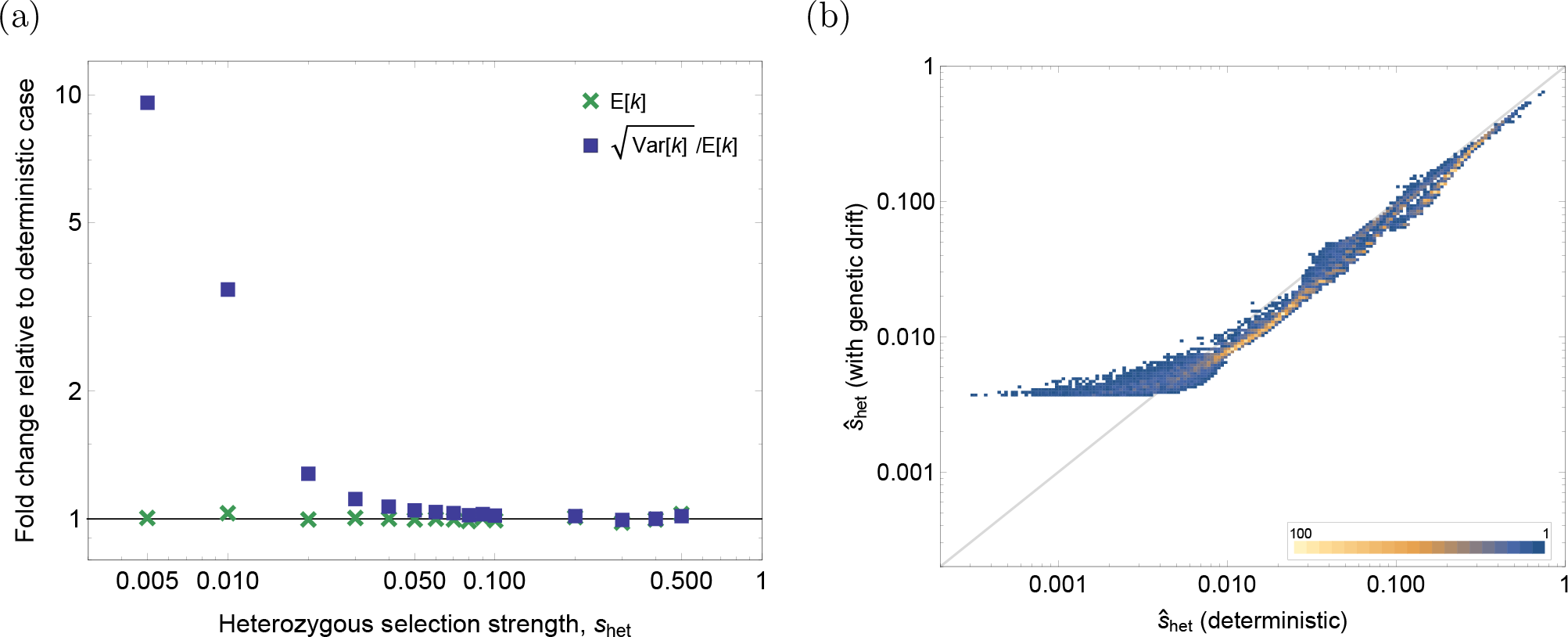
Comparison of the deterministic mutation-selection balance model with the model that includes the effects of genetic drift, in the NFE demography. (a) Fold-change in the coefficient of variation (squares) and the mean (crosses) of the number of PTV mutations, *k*, relative to the deterministic case, obtained from simulations of a realistic demography of the ExAC NFE sample for different values of heterozygous selection strength *s*_het_. (b) Heat map of gene-specific estimates for all 16,279 used genes from the NFE sample, showing deterministic (x-axis) and drift-inclusive (y-axis) *s*_het_ estimates. Note the double-logarithmic axes in both panels.

## Incorporation of genetic drift in the selection inference

Having established the effects of genetic drift on the population PTV allele frequency, we could then address the resulting effects on selection inference. Based on the approach described in ref. [1], we estimated the parameters ***θ*** of the distribution of heterozygous selection coefficients, *P*(*s*_het_; ***θ***), from fitting the observed distribution of per-gene PTV counts,

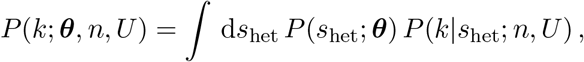

where *P*(*k*|*s*_het_; *n*, *U*) denotes the likelihood function of count *k* given *s*_het_. At mutation-selection balance, *P*(*k*|*s*_het_; *n*, *U*) = Pois(*k*; *nU*/*s*_het_), with Pois(*k*; *λ*) denoting the Poisson distribution of *k* with parameter *λ*.

Here, we fully accounted for the effects of drift through incorporation of the allele frequency spectrum, *ϕ*(*X*; *s*_het_, *U*). In the deterministic case, *X* is assumed to be fixed at its expected value *U*/*s*_het_ for a gene with mutation rate *U* and under selection *s*_het_. In contrast, we now explicitly include its variability:

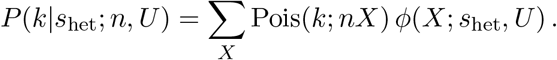

Population allele frequency spectra *ϕ*(*X*; *s*_het_, *U*) under the described demography were produced from simulations for a dense grid of mutation rates and selection strengths (**Methods**). We maximized the likelihood *P*(*k*; ***θ***, *n*, *U*) over parameters ***θ***, fitting this model to the observed distribution of *k* in the ExAC NFE subset (**Supplementary Figure 1**). As we found previously, the inverse Gaussian distribution provided the best parametric form for the fit [1]. To obtain gene-specific estimates of selection, *ŝ*_het_, we derived the mean of the posterior distribution (**Methods**).

We used the resulting per-gene estimates to understand the impact of a full demographic model relative to the deterministic estimates of heterozygous selection coefficients obtained under mutation-selection balance. **Figure 1(b)** shows the direct comparison between the two scenarios for the NFE subset. Overall, the incorporation of drift in the model did not substantially change relative ranks of genes (Spearman rank correlation coefficient = 0.995; **Supplementary Figure 2**). Estimates for genes under moderate to strong selection (*s*_het_ > 0.01, N=10,744 genes, 66%) are very close to the deterministic estimates, only showing a slight downward shift on average. As variance due to genetic drift partly absorbs the variance of the prior distribution of selection coefficients, the latter decreased (by about 18%). Genes under weaker selection (*s*_het_ < 0.01, N=5,535 genes, 34%) appeared with largely the same rank, but showed a monotonic increase in estimated values of heterozygous selection. The apparent convergence of selection coefficients to ~0.004 for these genes can be explained by the effects of genetic drift on the likelihood function. For weak negative selection, *P*(*k*|*s*_het_; *n*, *U*) becomes nearly flat in the regime of very small *s*_het_, moving the estimates closer to the genome-wide mean.

We conclude that even though human demographic history is complex, a realistic model of recent population expansion suggests that, owing to their deleteriousness, the evolution of PTVs can be largely described in a deterministic framework.

## Test for differences between subpopulations

As an additional way to test the utility and limits of the deterministic approximation to estimate selection, we used a data-driven approach. This approach relied on a comparison of PTV counts between different human subpopulations, free from assumptions made in simulations, such as panmixia. For a given gene, if the same selection coefficient *s*_het_ acts on heterozygous PTVs in two subpopulations, we expect E[*X*_1_] = E[*X*_2_] = *U*/*s*_het_. Here *X_i_* denotes the cumulative PTV allele frequency in subpopulation *i*. The PTV allele count in subsample *i* was modeled as *k_i_* ~ Pois(*n_i_U*/*s*_het_), where *n_i_* is the number of chromosomes sampled.

We tested whether this deterministic approximation was violated by comparing the Non-Finnish European (NFE, *i* =1) and African (AFR, *i* =2) ExAC subsamples using the C-test [17]. Given *k*_1_ + *k*_2_ ≡ *k*, the distribution of *k*_1_ conditional on the total count *k* is binomial with success probability *p* = *n*_1_/(*n*_1_ + *n*_2_). We then computed the two-sided binomial p-value of the observed value of *k*_1_ for all genes in the set. Since the *k*_1_ are discrete random variables, their p-values are not uniformly distributed. Therefore, in order to account for multiple testing, we compared the observed genome-wide distribution of binomial p-values to the p-value distribution obtained from a simulation under the null assumption. We generated 500 instances of simulated binomial PTV counts for each gene and computed the false discovery rate (FDR) conservatively, as the fraction of genes that is expected by chance to have a p-value equal to or less than a certain threshold. We then measured the number of genes with FDR below 5%, *x*_sig_.

**Table 1** shows the results for different intervals of the deterministic *s*_het_ estimates, as well as the total number of genes in each interval, *x*_tot_, and significant fraction. We found that a total of 870 out of 15,865 used genes (5.5%) show a significant deviation from the assumption of Poisson distributed counts with equal expected values in the two subpopulations. Of all 870 significant genes, 49% have *ŝ*_het_ ≤ 0.01. Unlike our simulations, this data-driven approach is not expected to be contingent on assumptions about demographic structure or aspects of the population history.

**Table 1.**
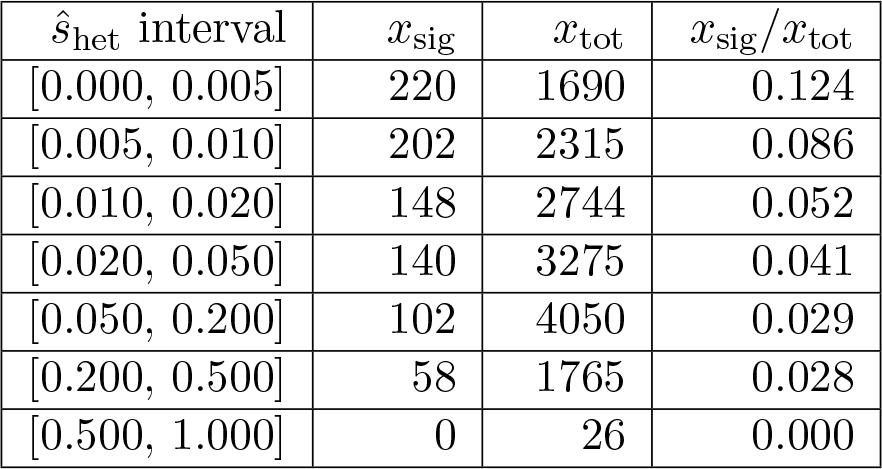
Fraction, *x*_sig_*/x*_tot_, of genes with significant, FDR-corrected two-sided binomial p-value according to the C-test across the NFE and AFR subpopulations (out of N=15,865 genes), in intervals of deterministic *ŝ*_het_ values derived from the NFE subpopulation of the ExAC dataset. FDR was controlled at 0.05.

## Prediction of *de novo* fraction using *ŝ*_het_

Some of the PTVs detected in a sample are *de novo* mutations rather than segregating alleles inherited from the parental generation. With increasing strength of negative selection, the population allele frequency, and thus the chance of inheriting a deleterious allele, is reduced and more of the observed deleterious mutations arise *de novo*. The fraction of *de novo* out of all PTVs equals *s*_het_ for genes under negative selection in the deterministic limit. As shown in **Methods**, this result is also valid across a wide range of parameters at mutation-selection-drift balance.

We have collected *de novo* and inherited PTVs in autism-spectrum disorder probands from 3,009 parent-child trios [18]. For each gene, we computed the observed fraction of de novo PTVs, 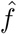, and compared it to the deterministic estimate of the heterozygous selection coefficient, *ŝ*_het_. This analysis provides another independent and data-driven approach to test the validity of the *s*_het_ estimates. **Figure 2** shows the observed relation between *ŝ*_het_ and 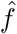. We find very good agreement between *ŝ*_het_ and 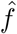 in the limit of intermediate to strong selection (*ŝ*_het_ ≥ 0.02). Genes with smaller *ŝ*_het_ largely show agreement with 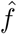 within error bars. **Supplementary Figure 3** shows the corresponding plot for the deterministic *s*_het_estimates obtained from the entire ExAC dataset [1].

**Figure 2:**
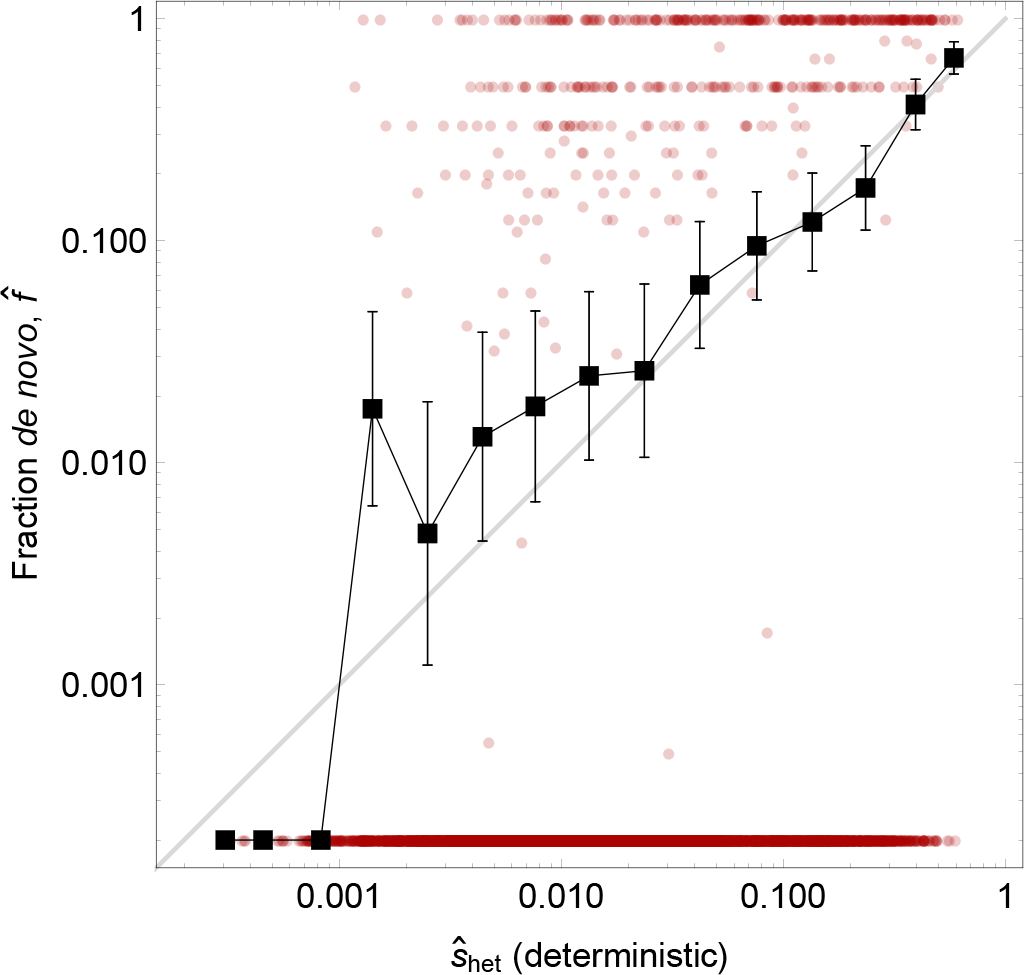
In the strong selection limit, *s*_het_ is a predictor of the fraction of *de novo* mutations *f* (**Methods**). *De novo* fraction 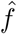 of PTV mutations was computed for 7,105 (out of 16,279) genes with at least one PTV (*de novo* or transmitted) in an autism-spectrum disorder cohort of 3,009 parent-child trios [18] (y-axis) and compared to the deterministic *ŝ*_het_ derived from the NFE sample (x-axis). Red dots denote individual genes (genes with 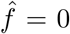 were assigned *ŝ*_het_ = 2 · 10*−4* for illustration purposes). Black squares connected by black lines denote the mean in bins along the x-axis of logarithmic width Δlog[*ŝ*_het_] = 0.25 (number of genes per bin from left to right: {1, 13, 55, 168, 435, 877, 1282, 1300, 975, 703, 429, 443, 303, 117, 4}). Vertical and horizontal error bars show the standard error of the mean per bin for 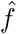 and *ŝ*_het_, respectively. Grey line denotes the diagonal.

## Comparison of *ŝ*_het_ to other measures of protein constraint

Beyond the theoretical importance, evaluating selection on deleterious PTVs has practical applications in human genetics. Population-based measures of constraint, such as pLI [2] and RVIS [19], have been successfully used to prioritize genes in studies of neuropsychiatric and other diseases [20]. pLI measures the probability of a gene to be loss-of-function intolerant [2]. This measure is based on a classification of genes into three categories and returns high values for genes under strong selection. By construction, this approach has limited resolution within this class of genes. Point estimation of *s*_het_ characterizes the fitness loss beyond the binary classification of whether or not the gene is under constraint. Therefore, *ŝ*_het_ has an advantage as a proxy for penetrance, disease age of onset and severity. This is illustrated in **Figure 3**, together with a comparison of the predictive power of pLI of the fraction of *de novo* mutations (see also **Supplementary Figure 4**).

**Figure 3:**
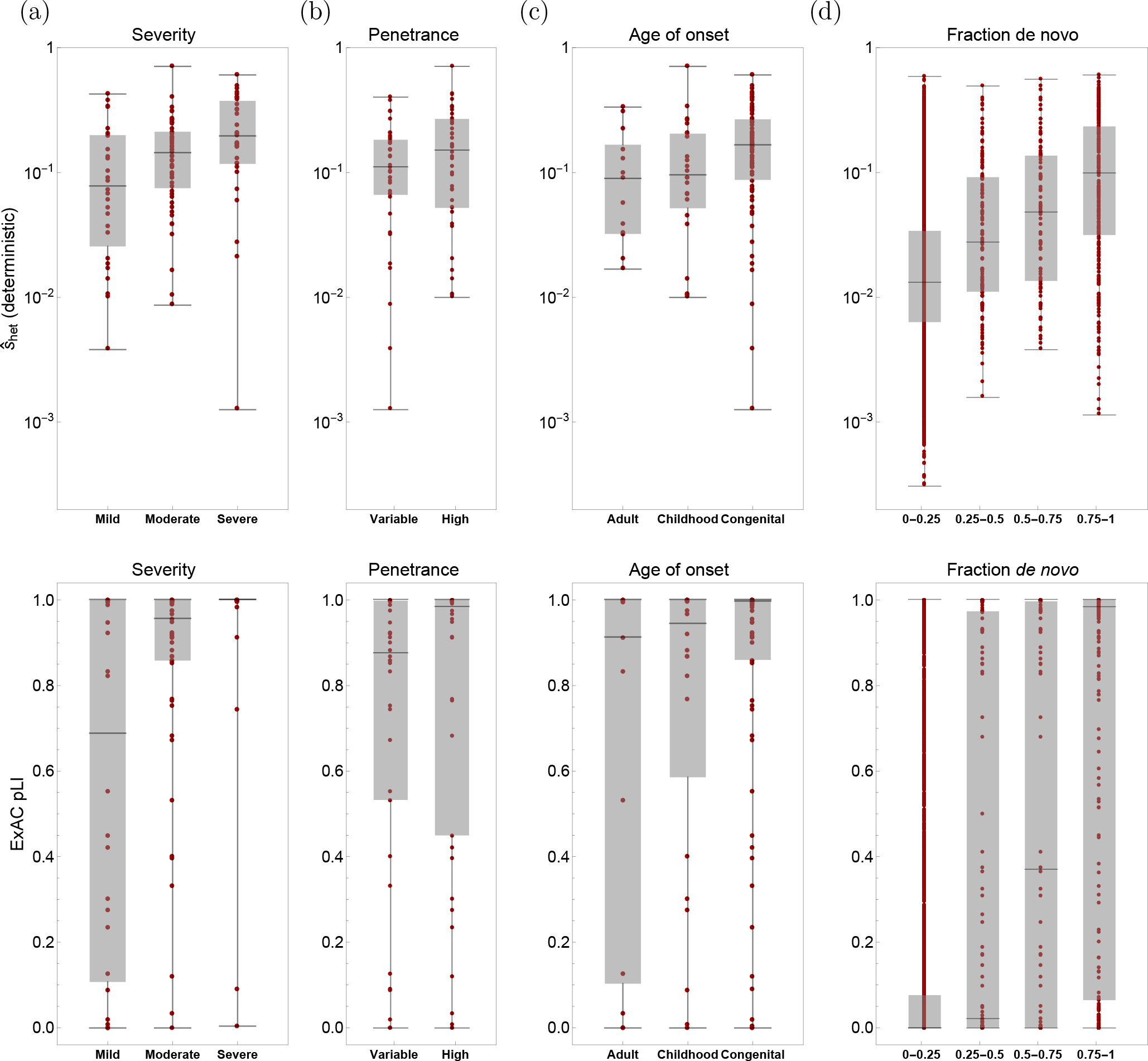
Comparison of per-gene selection estimates, *ŝ*_het_, with a measure of probability of loss-of-function intolerance, pLI [2]. Shown is the correlation with independent measures of gene importance. Red dots denote individual genes, grey boxes enclose the central quartiles of the distribution in each category, and black horizontal bars through grey boxes show the median. (a-c) Data on disease severity, penetrance, and age of onset (x-axes) for a set of 112 haploinsufficient disease-associated genes of high confidence (ClinGen Dosage Sensitivity Project) were compared to deterministic *ŝ*_het_ from the NFE sample (top row) and pLI (bottom row) predictions (y-axis). (d) The fraction of *de novo* PTVs, shown for 6,811 genes in bins of width 0.25 on the x-axis, was derived from a trio-sequencing dataset [18] (top: *ŝ*_het_, bottom: pLI). Note the logarithmic y-axis in the top row, while the bottom row has a linear y-axis.

## Conclusions

The existence of stochastic effects in finite populations has long been known [4], predating diffusion theory approaches to the evolutionary dynamics of populations [7]. However, the impact of genetic drift on the fate of a newly arising mutation was not always appreciated. Partially due to historically insufficient amounts of data, the deterministic forces of mutation and selection were often the focus of population genetics analyses, corresponding to working in the limit of infinite effective population size, *N*_e_. The equilibrium state in that case is mutation-selection balance. Arguably, a locus evolving under strong selection and high mutation rate may still be considered in mutation-selection balance even if the effective population size is finite (that is, if it is large enough to ensure 4*N*_e_*U* ≳ 1 and 4*N*_e_*s*_het_ > 1). When considering a gene as a biallelic locus that can either carry or not carry a variant under strong negative selection, the cumulative mutation rate to the deleterious state is sufficiently large in human populations to satisfy this limit. In our earlier work, we hypothesized that also selection against heterozygous protein-truncating variants scaled by *N*_e_ would be sufficiently strong to ensure the assumption of mutation-selection balance [1].

Here we have investigated the full effects of drift using simulations of a realistic human demography. This approach models recurrent mutations, is gauged by the observed NFE data in the ExAC cohort, and does not make assumptions about equilibrium. As a result of our analysis, we found that the deterministic mutation-selection balance approximation for counts of rare deleterious PTVs is applicable for genes under strong to intermediate selection, including genes with the previously inferred global mean of *ŝ*_het_ ≈ 0.06. Selection estimates for this class of genes are highly robust to the incorporation of drift effects. For genes under relatively weak selection, the deterministic *s*_het_ estimates provide a stable ranking useful as prioritization scores for practical applications in human genetics [1].

The ExAC dataset used to estimate the deterministic *s*_het_ given in ref. [1] is composed of different subpopulations. Here we focused on the largest subset, NFE, with its established demographic dynamics, to compare deterministic evolution of PTVs to a scenario under genetic drift. Population structure can cause an increase in the variance of allele frequency. In the case of lethal but highly recessive variants, the mean allele frequency can even be reduced below the deterministic expectation of the combined population [21, 11, 12]. To address the effects of population structure on the deterministic *s*_het_ estimates, we conducted two data-driven tests. First, we tested whether the PTV allele counts in the Non-Finnish European and the African ExAC subsamples were compatible with being generated under the same deterministic Poisson model with identical selection strength. Second, we analyzed the relationship between our estimates of heterozygous selection coefficients based on the combined ExAC dataset and the experimentally determined fractions of de novo mutations (**Supplementary Figure 3**). Both of these analyses are consistent with the simulation results, suggesting that the mutation-selection balance approximation is applicable to strongly to moderately selected genes.

## Acknowledgements

We kindly thank Brian Charlesworth and William Hill for their interest in our work and for inspiring this study. We would like to thank Jeremy Berg, Guy Sella, Molly Przeworski and Alexey Kondrashov for their helpful suggestions and feedback.

## Methods

### Analytical derivation of PTV count variance

We define the cumulative PTV allele frequency of a gene in the population, *X* = ∑_*j*_ *x*_*j*_, where the sum is over all PTV loci *j* on the gene with respective allele frequencies *x_j_*. The frequency *X* is governed by the demographic dynamics of the population. For heterozygous selection coefficient *s*_het_ and cumulative mutation rate *U*, it follows the allele frequency distribution *ϕ*(*X*; *s*_het_, *U*). Given frequency *X* in the population, we expect to see on average E[*k*] = *nX* mutations in a sample of *n* chromosomes, hence

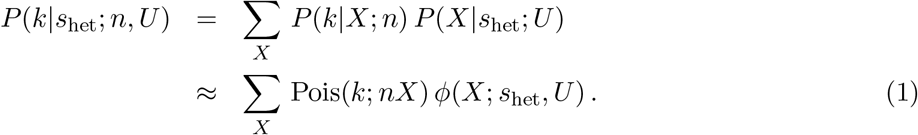

Here, Poisson sampling with parameter *nX* again represents the limit of binomial sampling for small success probabilities *x_j_* and large sample size *n*. In particular, we are interested in how genetic drift affects the mean and variance of the PTV count *k*. For a given selective effect *s*_het_ we obtain the expected value of *k* from Eq. 1,

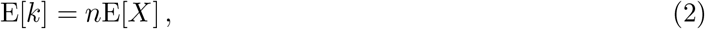

while the variance is given by

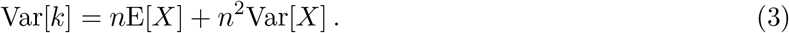

### Analytical approximation for strong selection on heterozygous sites

Both E[*X*] and Var[*X*] depend on the evolutionary dynamics of the population, but it is instructive to compare to the equilibrium case of strong purifying selection on heterozygous variation described by Nei [12]. Nei showed that in that case the theoretical allele frequency distribution can be approximated by a gamma distribution with shape parameter 4*N*_e_*U* and scale parameter 1/(4*N*_e_*s*_het_), where *N*_e_ again denotes the effective population size. Under this assumption, Eqs. 2 and 3 become

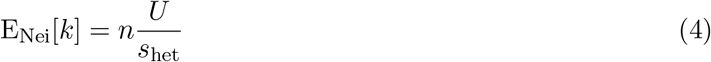

and

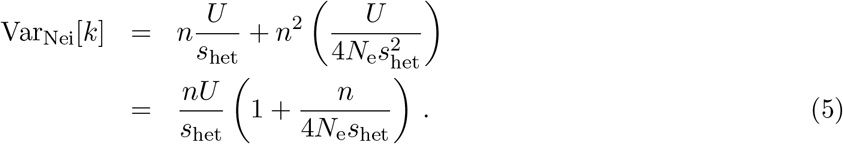

When we compare this to the analogous expressions for mean and variance under the deterministic assumption of mutation-selection balance,

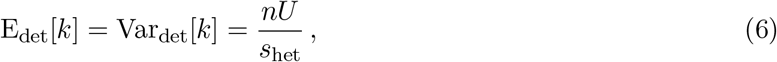

we find the same expected value. For the excess variance we consequently expect to see a suppression with increasing effective size of the population from which the sample was drawn, as well as with increasing selection strength.

### Population genetics simulations

To understand the interaction between mutation, selection, genetic drift, and population sampling, we wrote a custom Wright-Fisher simulator appropriate for the parameter regimes of interest. We assumed infinite recombination, such that all sites evolve independently, in a panmictic population with a single value of genic (additive) natural selection per simulation, and a fixed mutation rate with no back mutations. We modeled genes as biallelic sites with a mutation rate associated with the sum of all PTV targets in the gene, *U*. For each simulated biallelic site, recurrent mutations were Poisson sampled with mean *AU*, where *A* is the number of ancestral, i.e. unmutated, chromosomes. For a given mutation rate and selection coefficient we generated 100,000 independent realizations, providing a distribution of biallelic frequencies. We simulated a dense grid of mutation rates (*U* ∈ [10^−7^, 7.16 × 10^−5^]) associated with the PTV-specific mutation rate estimates provided in Samocha *et al.* (2014) [22]. For each biallelic mutation rate, we simulated a range of 12 selection coefficients: *s*_het_ = {5 × 10^−6^, 1.6 × 10^−5^, 5 × 10^−5^, 1.6 × 10^−4^, 5 × 10^−4^, 1.6 × 10^−3^, 5 × 10^−3^, 1.6 × 10^−2^, 5 × 10^−2^, 1.6 × 10^−1^, 5 × 10^−1^, 1}.

We equilibrated simulations at the human ancestral population size of *N*_e_ = 14,474 for a burn-in period of 10*N*_e_ =144,740 generations. At this point, we simulated 5,921 generations of European demographic history, following the parameters specified in Tennessen *et al.* (2012) [9]. This corresponds to a constant population size for 3,880 generations, then a 1,120 generation bottleneck down to 2*N*_e_ = 3,722 individuals, then a 716 generation mild exponential growth phase with a growth rate of 0.00307, followed by a final exponential growth epoch of 205 generations. The final exponential growth phase was modeled three times with three distinct growth rates: growth = {0.0195, 0.024, 0.030}. The first two growth rates were chosen based on published values of the final European effective population size: 0.0195 from [9] and 0.024 roughly corresponding to Gao *et al.* (2014) [14], corresponding to final population sizes of around *N*_e_ = 0.5 million and *N*_e_ = 1.25 million, respectively. We estimated the final growth rate by simulating neutral variation through a dense range of growth rates and comparing to observations of the relative number of singletons to all segregating sites for synonymous sites in the ExAC NFE sample. This comparison was performed using a mutation rate of 3.8 × 10^−9^ for non-CpG transversions [23], and comparing to the fraction of non-CpG transversion singletons in the NFE sample. CpG and non-CpG transitions were explicitly ignored for these purposes, as they are known to exhibit the effects of recurrent mutations in the ExAC sample due to elevated mutation rates relative to non-CpG transversions. An effective population size growth rate of 0.03 per generation in the last exponential epoch provided a fraction of singletons of 0.636, which is highly consistent with the number of observed singletons in ExAC NFE of 0.633. Unsurprisingly, the final population size of *N*_e_ = 4.3 million found by matching the fraction of singletons is consistent with a recent population size inference performed on the same dataset [15].

We simulated all mutation rate and selection strength combinations through all three European demographies, and found a distribution of biallelic frequencies (including the number of monomorphic sites) for each. The distributions corresponding to the demography matching the ExAC NFE dataset (final growth rate of 0.03) were used in all subsequent analyses.

### Assessment of variance in comparison to Poisson sampling variance

To analyze the deviation from Poisson variance, we downsampled the final population of the simulation to the sample size of the NFE individuals in ExAC. The variance was computed over independent sites and compared to the theoretical mean of the Poisson distribution to assess the effects of genetic drift on the variance for a gene relative to the Poisson variance expected at mutation-selection balance.

**Figure 1(a)** shows the deviation of the coefficient of variation from the Poisson expectation due to the effects of a realistic demographic history (growth rate of 0.030, consistent with ExAC NFE observations). Simulations are shown for the non-CpG transversion mutation rate of 3.8 × 10^−9^, as the inflation of the Poisson assumption should be independent of the mutation rate. **Supplementary Figure 5** shows the same deviation in all three demographic models, suggesting that the slowest growth model [9] inferred from 1,351 European American exomes deviates at higher selection strengths than the demographic model associated with the much larger ExAC dataset. However, all three demographies are qualitatively consistent in their behavior, suggesting the departure from the Poisson assumption (variance to mean ratio >1.5) occurs at roughly *s*_het_ = 0.02, 0.03, and 0.06 for growth rates of 0.030, 0.024, and 0.0195, respectively.

### Hierarchical model to estimate *P*(*s*_het_; *θ*)

From Eq. 1 we derive the expression for the PTV count distribution when *s*_het_ is drawn from the distribution *P*(*s*_het_; ***θ***) with parameter vector ***θ***:

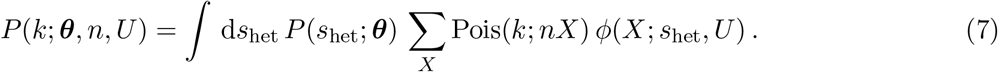

At mutation-selection balance, *X*_eq_ = *U*/*s*_het_ and *ϕ*(*X*; *s*_het_, *U*) = *δ*_*X,U*_/*s*_het_, where *δ* denotes the Kronecker delta, yielding

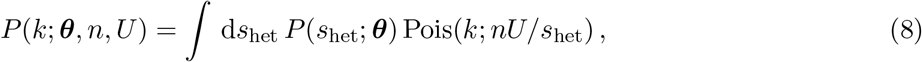

which exactly reproduces Eq. 4 of ref. [1].

### Fit to the PTV count distribution using simulations of a realistic demography

To evaluate the differences between the selection inference under the original mutation-selection balance assumption, Eq. 8, and the full model incorporating genetic drift, we applied Eq. 7 to fit the observed distribution of per-gene PTV counts. Because of the complex demographic histories of the different subpopulations constituting the ExAC dataset, particularly substantial recent population expansions, the simple equilibrium approximation by Nei [12] is not sufficient to substitute for *ϕ*(*X*; *s*_het_, *U*). Instead, we used the population allele frequency spectrum obtained from forward simulations described above, based on published demographies for the largest subpopulation, Non-Finnish Europeans. Since *ϕ*(*X*; *s*_het_, *U*) depends on *s*_het_, we used the simulated allele frequency spectra for each of 12 discrete values spanning six orders of magnitude (*s*_het_ ∈ [5 × 10^−6^, 1]) and rewrote the integral Eq. 7 as a sum. Genes were binned by their cumulative mutation rate *U* and for each *s*_het_ we simulated *ϕ*(*X*; *s*_het_, *U*) for *U*-bins of width 10^−7^ if *U* < 4 × 10^−6^, of width 2 × 10^−7^ for 4 × 10^−6^ < *U* < 1.4 × 10^−5^ and for exact *U* values above. Each spectrum contained 100,000 random instances of simulated frequencies *X*.

Parameters of the discretized and renormalized *P*(*s*_het_; ***θ***) were estimated from maximum likelihood. We compared the fits from three different two-parameter functional forms for *P*(*s*_het_; ***θ***): inverse Gaussian, inverse gamma, and gamma distribution, and found that the highest likelihood model is the inverse Gaussian, as in the deterministic case. Because the tested distributions are defined on the positive real axis, but the heterozygous selection coefficient *s*_het_ ≤ 1, *P*(*s*_het_; ***θ***) was truncated at *s*_max_ = 2.5, corresponding to a probability density of < 0.01. We find a slight dependence of the inferred parameters 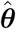 on 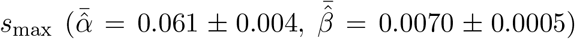, while the rank correlation changes by less than 0.1%. **Supplementary Figure 1** shows the resulting fit, 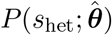, in comparison to the fit obtained under the mutation-selection balance assumption. As expected, the incorporation of genetic drift entails a moderate shrinkage of the variance (by 18%), as under the deterministic assumption its effects on PTV count variance are partly absorbed by 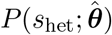. Per-gene estimates were derived as the mean of the posterior distribution:

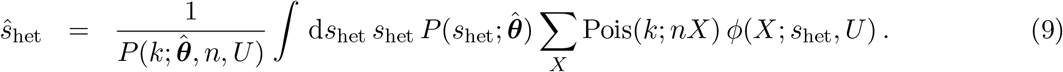

### Relationship between *s*_het_ and the de novo fraction of PTVs

In the limit of strong selection and at mutation-selection balance, the expectation for the cumulative PTV population allele frequency on a gene is *U*/*s*_het_. The relative fitness of an individual with a segregating PTV mutation in the current generation is 1 − *s*_het_, rendering the probability for a transmitted PTV in the next generation

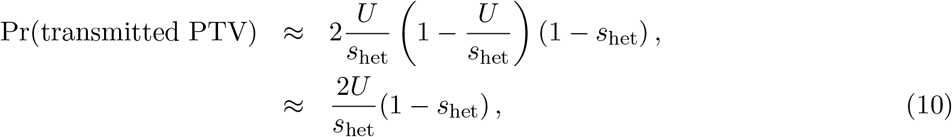

where we assumed *U* ≲ 10^−6^ ≪ 1 and the strong selection approximation *U*/*s*_het_ < 10^−3^ ≪ 1. The probability for a *de novo* PTV mutation to occur is

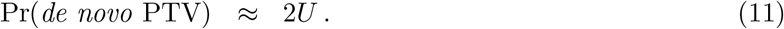

From this, we compute the fraction of *de novo* among all PTVs in the limit of strong selection:

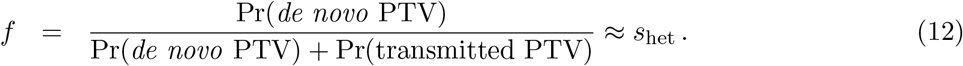

More generally, we can derive the fraction of *de novo* mutations from the equilibrium allele frequency spectrum for any value of *s*_het_. At mutation-selection-drift balance and in the limit of small mutation rates, the derived allele frequency spectrum is approximately given by

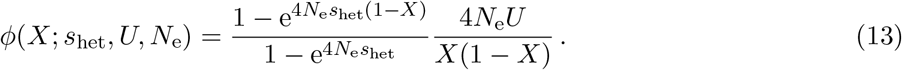

Hence, we obtain for the probability of inheritance

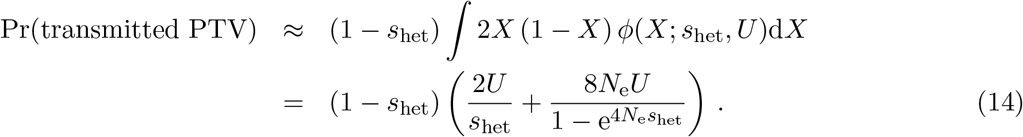

**Supplementary Figure 5** shows the resulting expression for the *de novo* fraction as a function of *s*_het_. Based on our estimates of recent effective population size, which is the most relevant for rare deleterious mutations, on the order of millions, we expect the one-to-one correlation between *s*_het_ and *f* to hold across the largest part of the selection regime. This renders the *de novo* fraction of PTV alleles an excellent cross-check for the *s*_het_ estimates. We computed the *de novo* fraction from an autism-spectrum disorder cohort [18], and all transmitted variants with genotype coverage ≥ 90% were included.

**Supplementary Figure 1:**
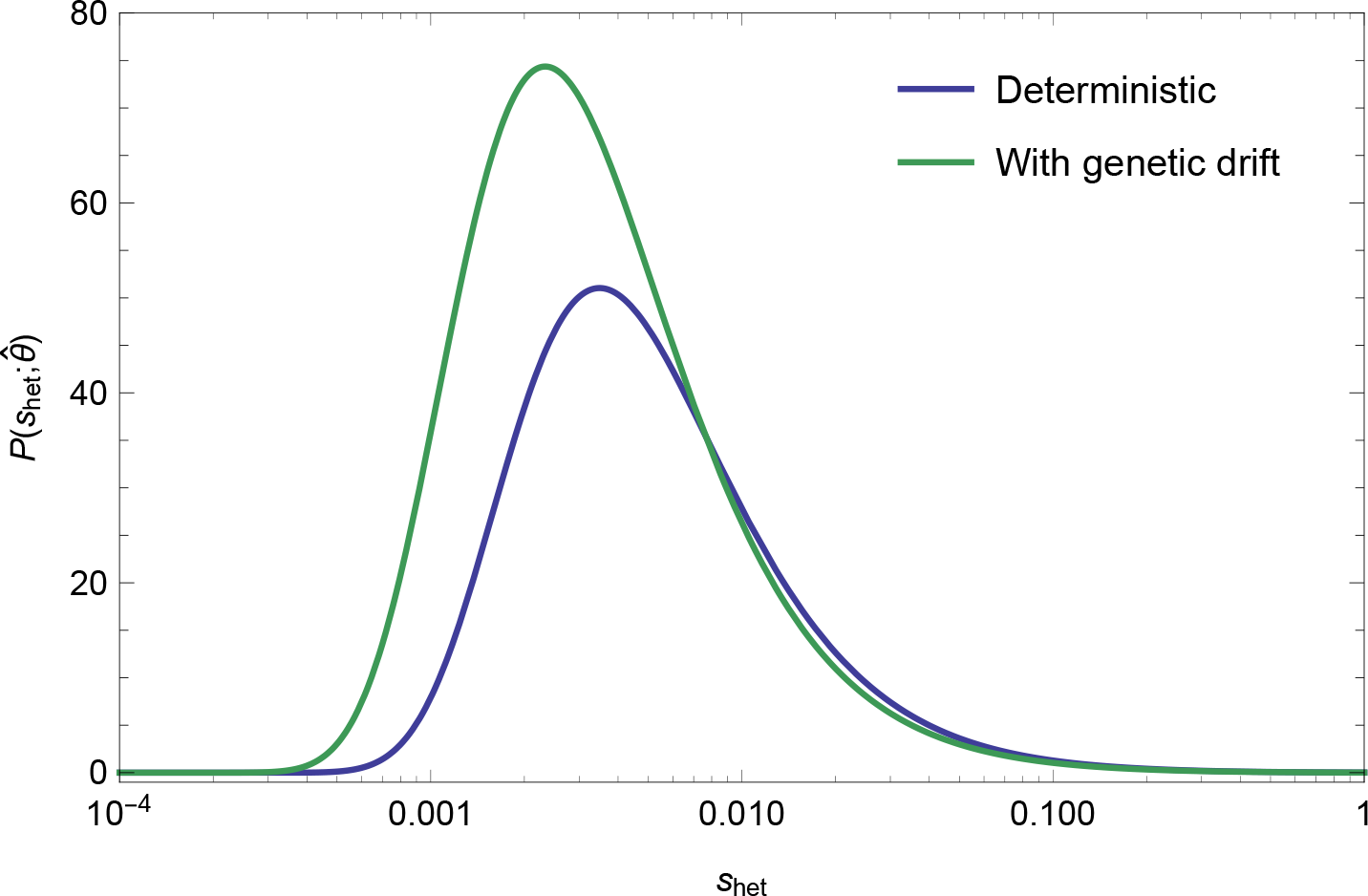
Comparison between 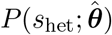 inferred under the mutation-selection balance assumption of Eq. 8 (blue) and when incorporating the effects of genetic drift, Eq. 7 (green). The latter scenario is based on a demographic model that fits the NFE subpopulation of the ExAC dataset and is in approximate correspondence with [15]. Distributions are shown for fits to the PTV counts in the NFE sample, containing 16,279 genes with cumulative PTV allele frequency 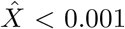. Estimated parameters are 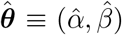, where *α* denotes the mean and *β* the shape parameter of the inverse Gaussian distribution. For the deterministic scenario we find 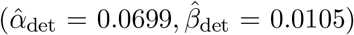, while including drift delivers 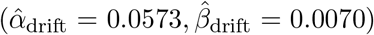, implying that the variance decreases by about 18% when stochasticity is accounted for. Note the logarithmic x-axis.

**Supplementary Figure 2:**
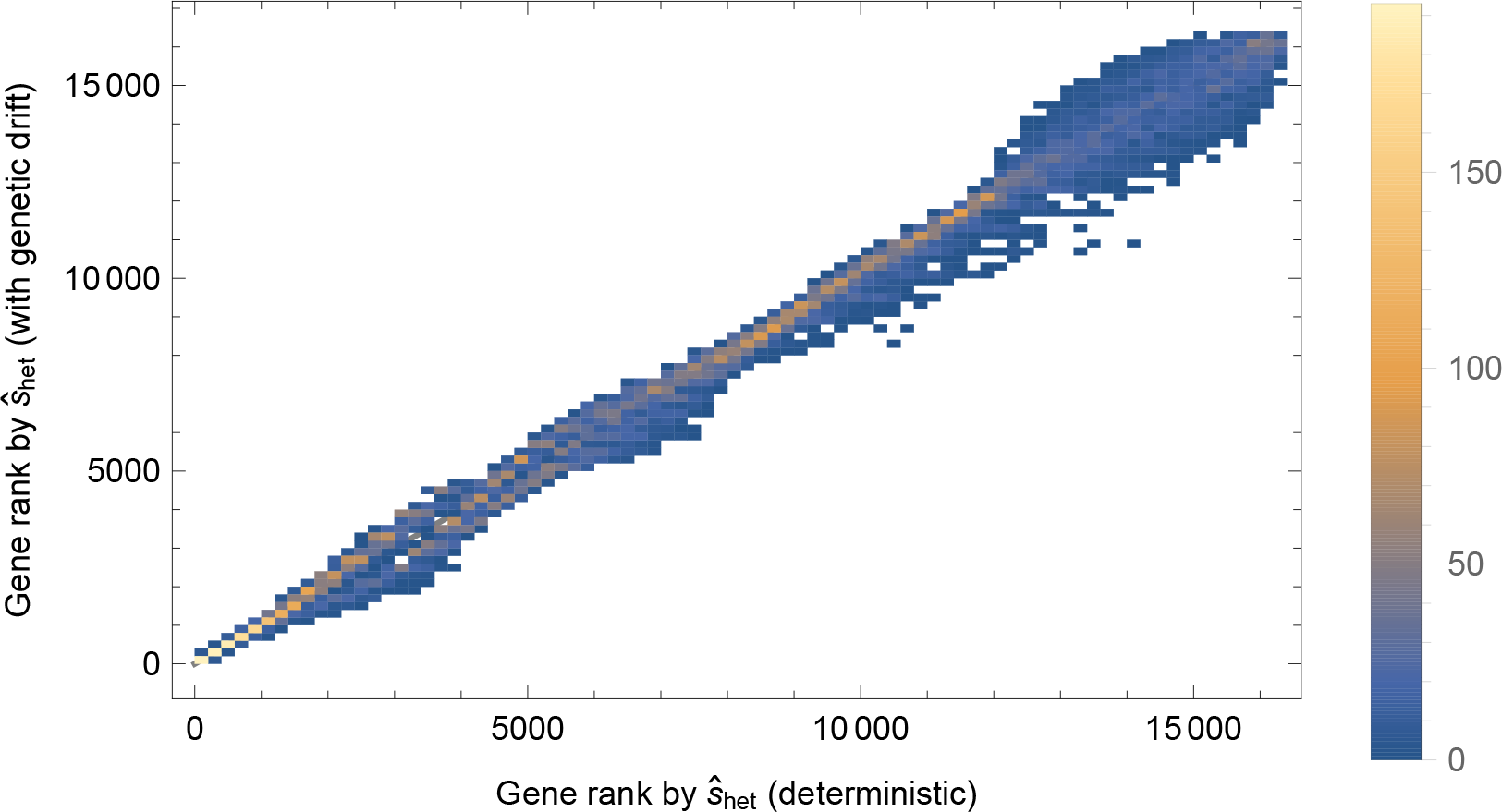
Comparison between the ranks of per-gene selection estimates *ŝ*_het_ for the NFE sample under the deterministic mutation-selection balance assumption (x-axis) and when incorporating the effects of genetic drift (y-axis). Shown is the density histogram from 16,279 genes. Spearman rank correlation between the estimates is 0.995.

**Supplementary Figure 3:**
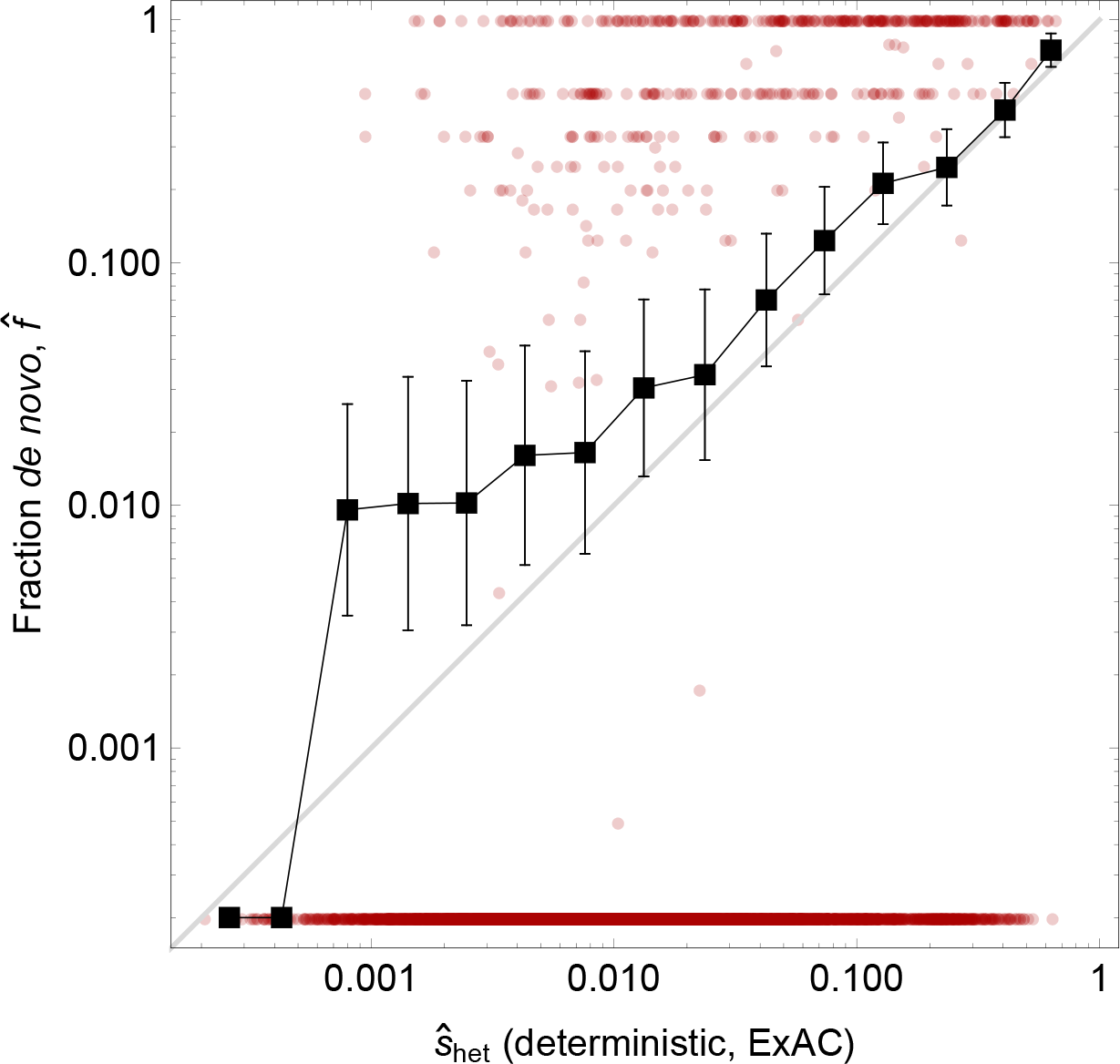
*De novo* fraction 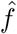 of PTV mutations was computed for 6,881 (out of 15,998) genes with at least one PTV (*de novo* or transmitted) in an autism-spectrum disorder cohort of 3,009 parent-child trios [18] (y-axis) and compared to the deterministic *ŝ*_het_ derived from the full ExAC sample [1] (x-axis). Red dots denote individual genes (genes with 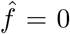 were assigned *ŝ*_het_ = 2 · 10^−4^ for illustration purposes). Black squares connected by black lines denote the mean in bins along the x-axis of logarithmic width Δlog[*ŝ*_het_] = 0.25 (number of genes per bin from left to right: {5, 31, 87, 295, 622, 1127, 1330, 1117, 809, 566, 358, 235, 217, 78, 4}). Vertical and horizontal error bars show the standard error of the mean per bin for 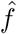 and *ŝ*_het_, respectively. Grey line denotes the diagonal.

**Supplementary Figure 4:**
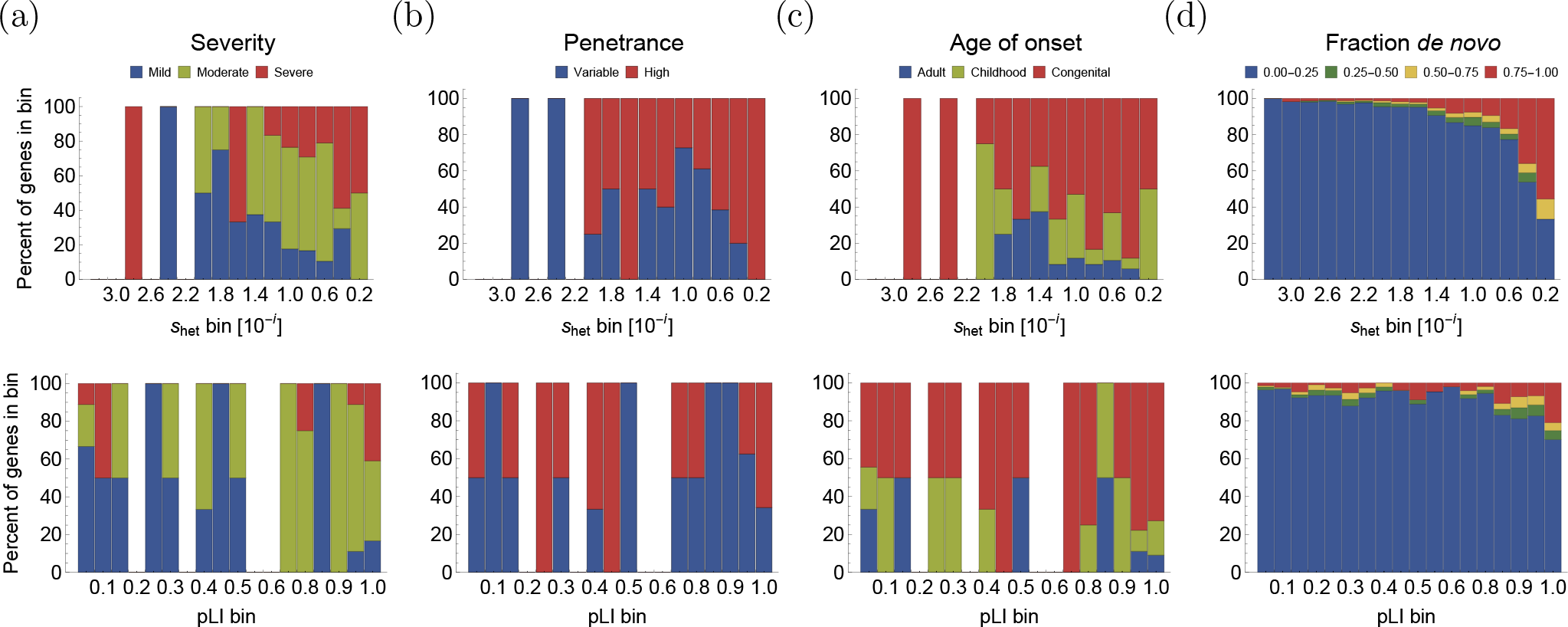
Comparison of per-gene selection estimates, *ŝ*_het_, with a measure of probability of loss-of-function intolerance, pLI [2]. Shown is a different depiction of the respective correlation of *ŝ*_het_ and pLI with independent measures of gene importance from **Figure 3**. Genes are binned by their *ŝ*_het_ values (top row, logarithmic bins of width 0.2) or pLI values (bottom row, linear bins of width 0.06) (x-axes). The y-axis shows the relative enrichment of each bin with the different categories of gene importance from each measure. (a-c) Data on disease severity, penetrance, and age of onset for a set of haploinsufficient disease-associated genes of high confidence (ClinGen Dosage Sensitivity Project). (d) The fraction of *de novo* PTVs, categorized by value into four bins of equal size, was derived from a trio-sequencing dataset [18]. The gene set used for severity, penetrance and age of onset is enriched for high *ŝ*_het_, reflected by the scarcity of observations in the low-*ŝ*_het_ regime (*ŝ*_het_ < 10^−2^) (a-c, top row). Most pLI values are located in the first and last bin (pLI < 0.06 and pLI > 0.94), causing depletion in the intermediate pLI regime. Note the logarithmic x-axis in the top row, while the bottom row has a linear x-axis.

**Supplementary Figure 5:**
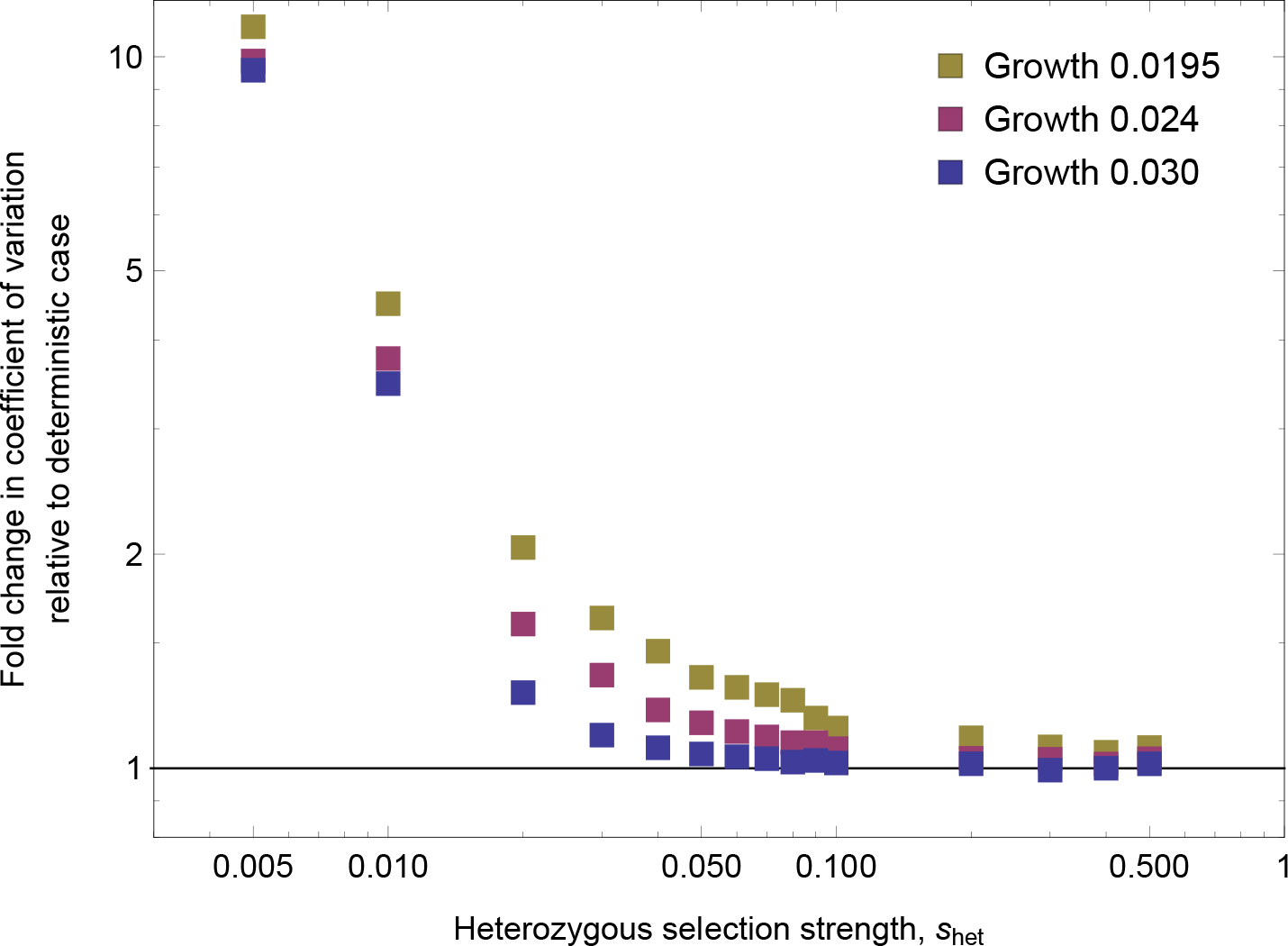
Fold-change in the coefficient of variation compared to the expectation under mutation-selection balance for different NFE demographies. Results are shown for three European demographies that differ in the rate of exponential expansion in the most recent exponential growth epoch. Inclusion of the effects of drift suggests a deviation from the Poisson assumption (Var[*k*]/(*nU*/*s*_het_) > 1.5) at approximately *s*_het_ = 0.02, 0.03, and 0.06 for growth rates of 0.030, 0.024, and 0.0195, respectively. Simulated points are shown (in order from left to right) for *s*_het_ = {0.005, 0.01, 0.02, 0.03, 0.04, 0.05, 0.06, 0.07, 0.08, 0.09, 0.1, 0.2, 0.3, 0.4, 0.5}.

**Supplementary Figure 6:**
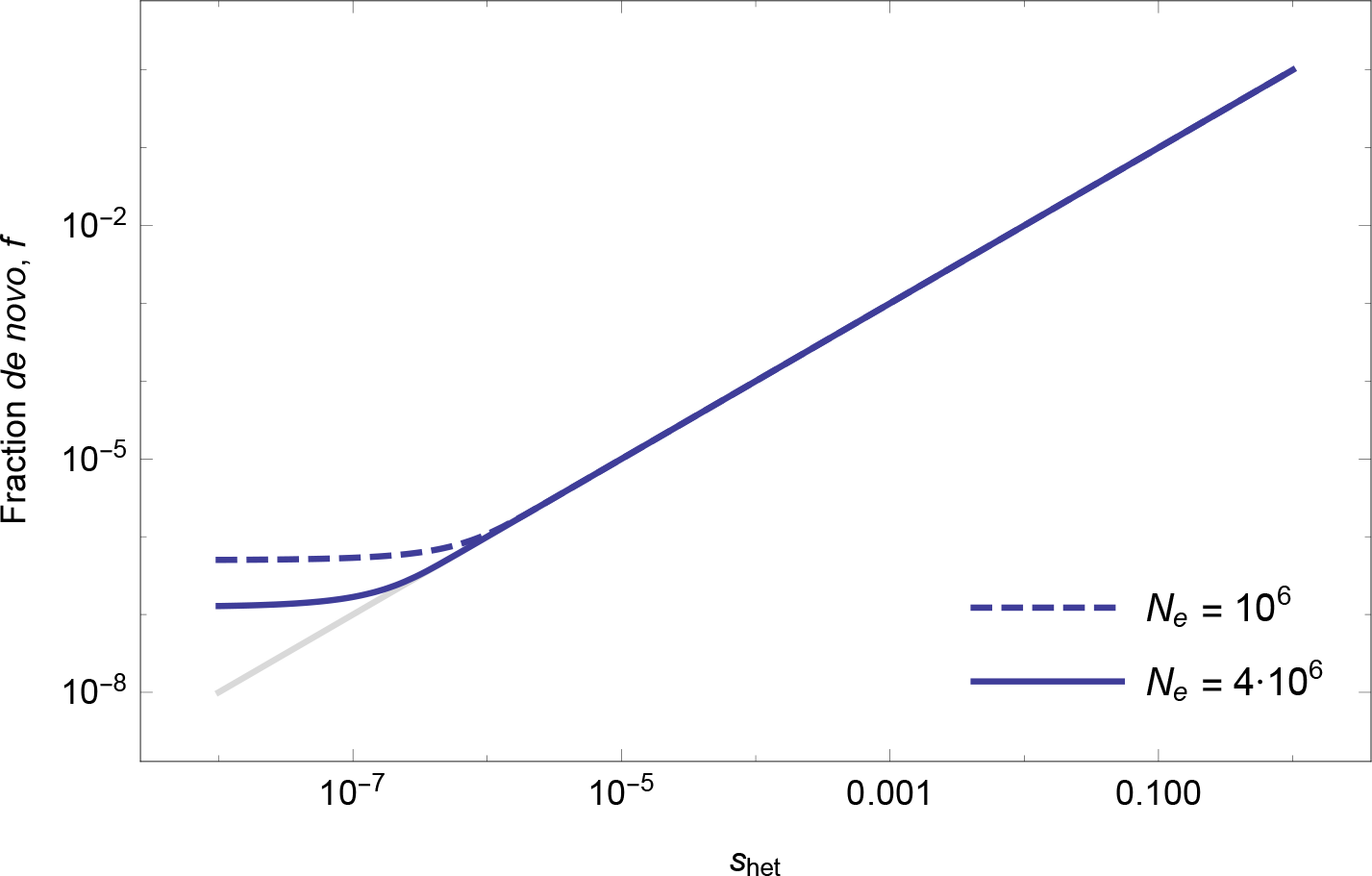
Analytical expression for the fraction of *de novo* mutations, *f*, occurring under heterozygous negative selection strength *s*_het_ in mutation-selection-drift balance for two different effective population sizes (solid blue line: *N*_e_ = 4 · 10^6^, dashed blue line: *N*_e_ = 10^6^). The grey line shows the diagonal. For very weak selection, *f* converges to the neutral equilibrium value of 1/(1 + 2*N*_e_).

